# 4-Hexylresorcinol Enhances Skeletal Muscle Glucose Handling through the AMPK-GLUT4 axis in Diabetic Rats

**DOI:** 10.64898/2026.01.19.700277

**Authors:** Xiangguo Che, Seung-Ki Hong, Ji-Hyeon Oh, Suyeon Park, Jongho Choi, Suk-Keun Lee, Dae-Won Kim, You-Young Jo, Seong-Gon Kim, Je-Yong Choi

**Affiliations:** Department of Biochemistry and Cell Biology, Cell and Matrix Research Institute, School of Medicine, Kyungpook National University, Daegu 41944, Republic of Korea; Department of Oral and Maxillofacial Surgery, College of Dentistry, Gangneung-Wonju National University, 7 Jukheon-gil, Gangneung-si, Gangwon-do 25457, Republic of Korea; Department of Oral Pathology, College of Dentistry, Gangneung-Wonju National University, 7 Jukheon-gil, Gangneung-si, Gangwon-do 25457, Republic of Korea; Institution of Hydrogen Magnetic Reaction Gene Regulation, Daejeon 34140, Republic of Korea; Department of Oral Biochemistry, College of Dentistry, Gangneung-Wonju National University, Gangneung 25457, Republic of Korea; Industrial Entomology Division, National Institute of Agricultural Science, RDA, Wanju 55365, Republic of Korea

**Keywords:** 4-hexylresorcinol, AMP-activated protein kinase (AMPK), GLUT4, skeletal muscle, insulin resistance, streptozotocin

## Abstract

**Background:** Skeletal muscle insulin resistance in diabetes is characterized by impaired GLUT4 availability and dysregulated AMP-activated protein kinase (AMPK) signaling. We investigated whether 4-hexylresorcinol (4HR) modulates glucose metabolism and AMPK–acetyl-CoA carboxylase (ACC)–GLUT4 signaling in vitro and in streptozotocin (STZ)–induced diabetic rats.

**Methods:** Differentiated C2C12 myotubes were treated with 4HR (0.1–10 µM) and assessed for glucose uptake, GLUT4 translocation, protein stability, proteasome involvement, and AMPK/ACC phosphorylation. In vivo, streptozotocin (STZ)–induced diabetic rats received subcutaneous 4HR (10 mg·kgL¹, 5 times/week for 7 weeks). Glycemic control, insulin secretion, muscle glycogen content, and serum protein profiles were evaluated.

**Results:** 4HR markedly increased glucose uptake (>5-fold vs. control, p<0.05), accompanied by increased phosphorylation of AMPK (Thr172) and ACC (Ser79), and redistribution of GLUT4 toward the cell periphery. Cycloheximide chase and proteasome inhibition experiments indicated partial stabilization of GLUT4. In diabetic rats, 4HR reduced fasting glycemia, improved glucose tolerance, elevated circulating insulin, and restored glycogen-rich fibers in skeletal muscle. Serum profiling revealed a metabolic and anti-inflammatory shift, with increased AMPKα, p-ACC, GLUT1, LGR4, LDHA, and PGC-1α, alongside suppression of NF-κB and TNF-α.

**Conclusion:** 4HR improves skeletal muscle glucose metabolism in STZ-induced diabetes, associated with activation of AMPK–ACC signaling, preservation of GLUT4 expression, and restoration of glycogen content. These findings highlight 4HR as a promising candidate for further investigation in diabetic muscle dysfunction and metabolic disease.

## 1. Introduction

Diabetes mellitus is globally prevalent and a chronic metabolic disorder characterized by persistent hyperglycemia and multisystem dysfunction [1]. Beyond its well-recognized microvascular and macrovascular complications, diabetes is increasingly recognized as a major driver of sarcopenia, the progressive decline of skeletal muscle mass and function [2,3]. Diabetic sarcopenia exacerbates frailty, reduces exercise capacity, impairs glucose disposal, and worsens clinical outcomes [4]. Epidemiologic studies consistently demonstrate reduced muscle mass and strength in individuals with diabetes [5], while experimental models recapitulate these deficits, underscoring a bidirectional relationship between diabetes and muscle dysfunction [6]. Diabetes accelerates muscle atrophy and impairs muscle function, further compromising glucose clearance, highlighting skeletal muscle as a central determinant of postprandial glucose uptake and future diabetes risk [2].

Mechanistically, diabetic sarcopenia is multifactorial. Insulin deficiency or resistance impairs GLUT4 translocation and glucose uptake, depleting glycogen stores and limiting ATP availability [7]. Catabolic signaling via FoxO transcription factors and the ubiquitin–proteasome system, together with dysregulated autophagy and mitochondrial dysfunction, compromise protein quality control and energy metabolism [8,9]. Inflammatory mediators and oxidative stress further disrupt excitation–contraction coupling and protein synthesis [10,11], collectively resulting in smaller myofibers, reduced contractile force, and early fatigue [12]. At the intersection of these pathways lies AMPK, a master energy sensor that facilitates GLUT4 trafficking and sustains muscle energetics [13,14].

Despite its clinical relevance, management of diabetic sarcopenia remains inadequate. Conventional glycemic control only partially restores muscle quality, and certain antidiabetic agents induce weight loss with disproportionate reductions in lean mass [15]. Nutritional supplementation (e.g., protein, leucine) and resistance exercise are beneficial but often unsustainable in chronic disease [16]. No approved pharmacotherapy exists, and investigational approaches such as myostatin/activin blockade or anabolic hormones have shown limited efficacy and raised safety concerns [17]. Thus, therapeutic agents that simultaneously enhance glucose handling and preserve muscle integrity represent a critical unmet need.

4-Hexylresorcinol (4HR) is a small phenolic compound with an established safety profile in food, dermatologic, and dental applications [18,19]. Emerging evidence suggests that 4HR exerts metabolic and cytoprotective effects, including AMPK activation [20], attenuation of oxidative and inflammatory stress [21], and modulation of proteostasis [22]. Given AMPK’s role in GLUT4 trafficking and muscle energetics [13,14], we hypothesized that 4HR might counteract key drivers of diabetic sarcopenia—namely impaired glucose uptake, glycogen depletion, and catabolic signaling. Supporting this rationale, our preliminary data in STZ-diabetic rat demonstrated that weekly 4HR treatment preserved muscle volume, restored PAS-detectable glycogen, and increased GLUT4 and p-AMPKα immunoreactivity in the masseter muscle [23].

In this study, we investigated whether 4HR attenuates diabetic sarcopenia and improves glycemic control in streptozotocin (STZ)–treated rats. Mechanistically, our focus was on the activation of AMPK, which subsequently phosphorylates and inhibits acetyl-CoA carboxylase (ACC), thereby promoting fatty acid oxidation and facilitating glucose uptake through GLUT4 translocation. This regulatory cascade is referred to as the AMPK-ACC-GLUT4 axis. Specifically, we (i) examined GLUT4 regulation at transcriptional (qRT-PCR), trafficking (confocal microscopy), and post-translational levels (cycloheximide chase, MG132, Bay-3827); (ii) evaluated in vivo outcomes including fasting glucose, intraperitoneal glucose tolerance test (IPGTT), longitudinal body weight, and inverted-screen performance; (iii) quantified muscle glycogen via PAS staining (mean intensity and proportion of high-glycogen fibers); (iv) assessed GLUT4, p-AMPK, and p-ACC expression in muscle tissue; (v) profiled serum proteins related to metabolism and inflammation using IP-HPLC; and (vi) investigated pancreatic islet morphology and serum insulin levels.

By addressing a critical knowledge gap, namely the absence of effective pharmacological strategies for diabetes, this study provides novel evidence that 4HR may serve as a dual-action agent, enhancing glucose metabolism while preserving muscle integrity. These findings hold translational relevance for the development of therapeutic approaches aimed at improving muscle health and metabolic outcomes in patients with diabetes

## 2. Materials and Methods

### 2.1 Cell culture and treatments

C2C12 myoblasts (ATCC CRL-1772; verified mycoplasma-free) were cultured in high-glucose Dulbecco’s Modified Eagle Medium (DMEM) supplemented with 10% fetal bovine serum and 1% penicillin–streptomycin at 37 °C in a humidified atmosphere containing 5% COL. Myogenic differentiation was induced by replacing the medium with DMEM containing 2% horse serum for 5–6 days until multinucleated myotubes were formed.

Differentiated myotubes were treated with 4-hexylresorcinol (4HR, 0.1–10 µM), insulin (100 nM), the AMPK inhibitor Bay-3827 (10 µM), or the proteasome inhibitor MG132 (10 µM). Vehicle controls received equivalent volumes of DMSO. Detailed procedures are provided in Expanded Materials and Methods.

### 2.2 Glucose uptake and GLUT4 trafficking assays

Glucose uptake was assessed using the fluorescent glucose analog 2-NBDG following serum deprivation and 4HR treatment. For GLUT4 trafficking, cells grown on coverslips were immunostained with anti-GLUT4 antibodies under permeabilized and non-permeabilized conditions, followed by confocal microscopy. Membrane-to-cytoplasmic fluorescence ratios were calculated from ≥20 cells per condition. Time-course studies, cycloheximide chase experiments, and MG132 cotreatment were used to evaluate GLUT4 synthesis and proteostasis. Methodological details are described in Expanded Materials and Methods.

### 2.3 Western blotting and quantitative RT-PCR

Protein lysates were analyzed using SDS-PAGE followed by immunoblotting for GLUT4, AMPKα, phospho-AMPKα (Thr172), and phospho-ACC (Ser79). Band intensities were normalized to β-actin. Total RNA was isolated for qRT-PCR to quantify GLUT4 mRNA using GAPDH as the reference gene. Primer sequences and complete protocols are provided in Expanded Materials and Methods.

### 2.4 Animal studies

Male Sprague–Dawley rats (n = 22; 6 weeks old) were used under an approved Gangneung-Wonju National University protocol (GWNU-2024-24). Diabetes was induced by intravenous streptozotocin (STZ, 50 mg·kgL¹) using a two-stage protocol. Rats with fasting glycemia > 300 mg/dL were randomized to diabetic vehicle or diabetic + 4HR (10 mg·kgL¹, subcutaneous, 5 times/week for 7 weeks; n = 9/group). Non-diabetic rats served as controls (n = 4). Body weight was monitored weekly, and gastrocnemius wet weight was measured at sacrifice. Induction, randomization, and blinding procedures are detailed in Expanded Materials and Methods.

### 2.5 Glycemic outcomes and functional testing

Fasting blood glucose was measured after 8 h fast. Intraperitoneal glucose tolerance tests (IPGTT) were performed using 1 g·kgL¹ glucose. Neuromuscular function was assessed by Kondziela’s inverted-screen test, and mean latency-to-fall from three trials was recorded.

### 2.6 Histological staining and immunohistochemistry

Gastrocnemius and pancreatic tissues were fixed, paraffin-embedded, and sectioned at 5 µm. Periodic acid–Schiff (PAS) staining was used to quantify muscle glycogen content, with high-glycogen fibers defined using an intensity threshold established from control samples. Immunohistochemistry (IHC) was performed for GLUT4, p-AMPK, and p-ACC. Antibody specifications and image-analysis procedures are provided in Expanded Materials and Methods.

### 2.7 Serum analyses

Plasma insulin concentrations were determined by ELISA. Serum metabolic and inflammatory proteins (including AMPKα, p-ACC, PGC-1α, GLUT1, NF-κB, TNF-α) were analyzed using immunoprecipitation-HPLC (IP-HPLC). Full analytic conditions are listed in Expanded Materials and Methods [24,25].

### 2.8 Statistical analysis

Data are presented as mean ± SD. Normality was assessed by Shapiro–Wilk test, and homogeneity of variance was evaluated by Levene’s test. Two-group comparisons were performed using Student’s t-tests, while comparisons among ≥3 groups were analyzed by one-way ANOVA followed by Tukey’s post hoc test. Time-course data were analyzed using two-way ANOVA with Geisser–Greenhouse correction and Sidak’s multiple comparisons test. Statistical analyses were conducted using GraphPad Prism v10, with significance set at α = 0.05.

## 3. Results

### 3.1 4HR enhances glucose uptake via GLUT4

To assess the effect of 4HR on cellular glucose uptake, differentiated myotubes were subjected to a fluorescent glucose analog assay (Fig. 2a). 4HR treatment significantly increased glucose uptake by 556.3% compared to the control and by 209.8 % versus the CHC-only groups (p < 0.05). Co-treatment with CHC, an inhibitor of clathrin-mediated endocytosis, markedly attenuated the 4HR-induced uptake, implicating a CHC-sensitive trafficking component. Insulin, used as a positive control, elicited the greatest uptake (p < 0.001). Confocal microscopy revealed weak perinuclear GLUT4 accumulation under basal conditions. Insulin increased perinuclear GLUT4 accumulation, whereas 4HR markedly enhanced overall GLUT4 expression. Notably, GLUT4 showed a more peripheral and diffuse cytoplasmic distribution compared with control, consistent with enhanced vesicle mobilization, although membrane fusion was not directly assessed (Fig. 2b).

**Figure 1.**
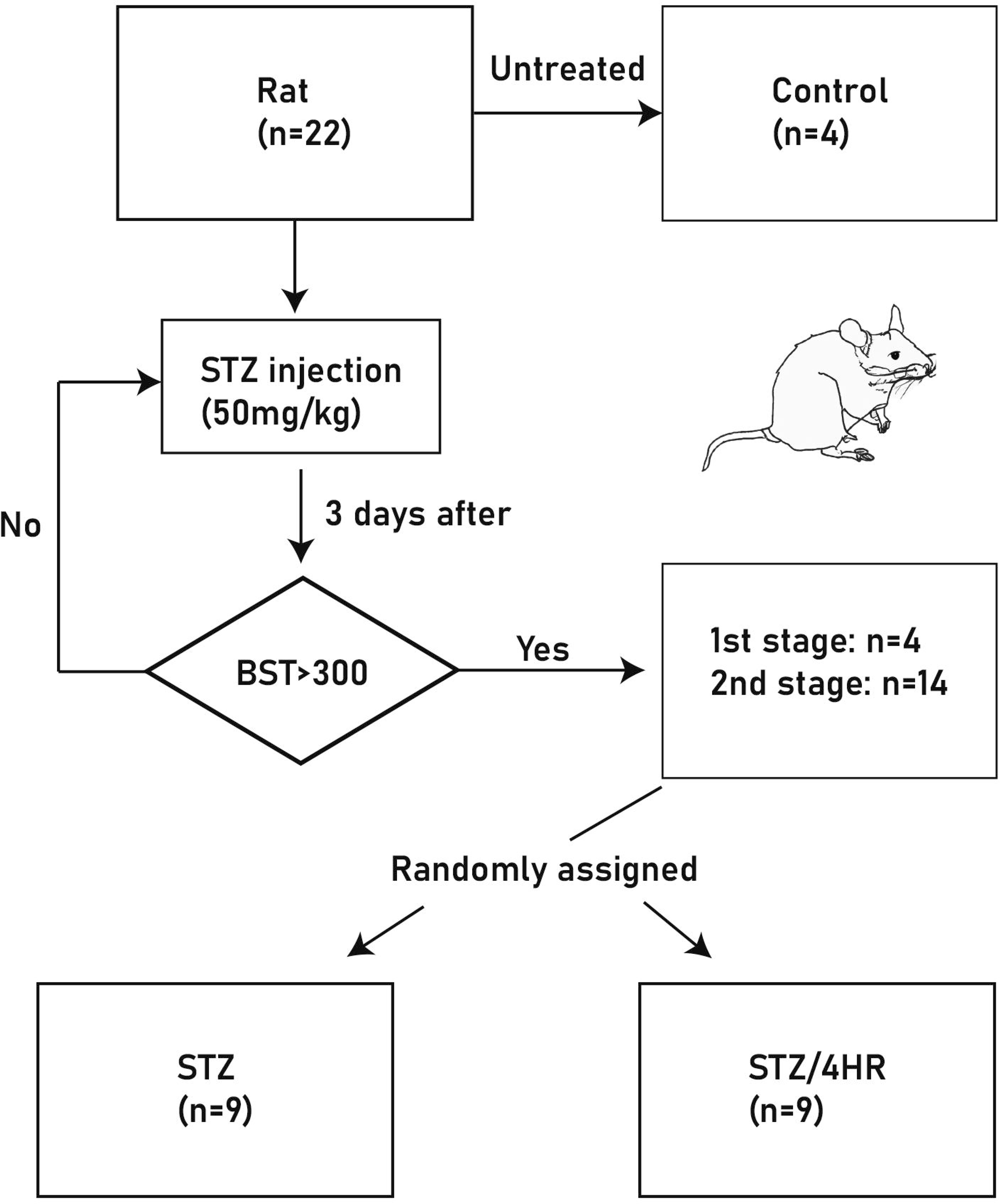
Experimental design of animal study. A total of 22 rats were used. Four rats served as the untreated controls (n=4). The remaining 18 rats received streptozotocin (STZ; 50 mg/kg) to induce diabetes. Three days after injection, blood sugar testing (BST) was performed. Rats with BST >300 mg/dL were considered diabetic and included in the study (stage 1: n=4; stage 2: n=14). Diabetic rats were then randomly allocated to either the STZ group (n=9) or the STZ/4HR treatment group (n=9).

**Figure 2.**
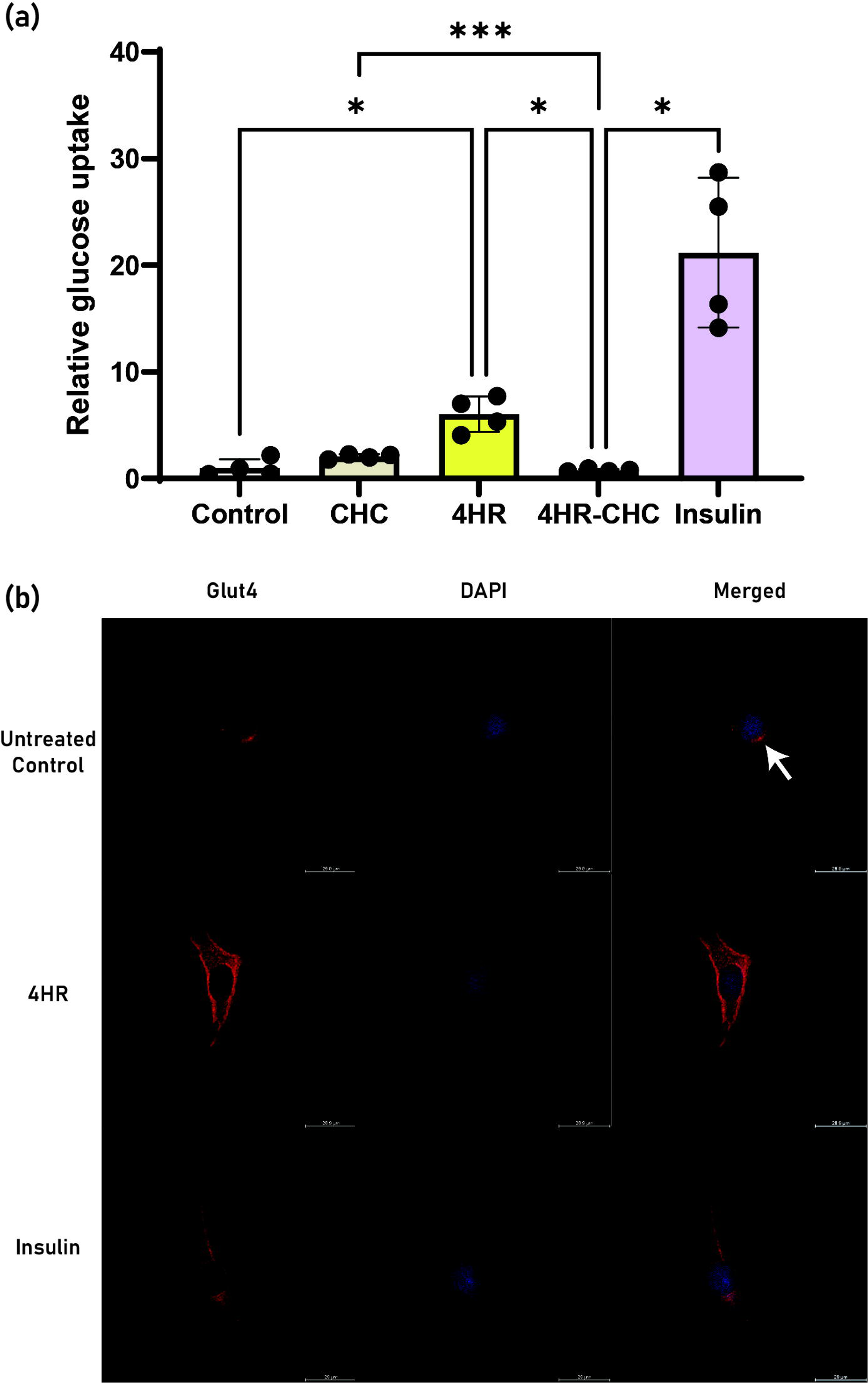
4HR enhances glucose uptake and GLUT4 redistribution. (a) Relative glucose uptake in C2C12 myotubes following treatment with CHC, 4HR, 4HR + CHC, or insulin. 4HR significantly increased uptake compared with control and CHC groups, whereas CHC cotreatment abolished this effect. Insulin elicited the highest uptake. Data are mean ± SD; *p < 0.05, ***p < 0.005. (b) Representative confocal images showing GLUT4 (red) and nuclei (DAPI, blue). Control cells exhibit perinuclear GLUT4 retention (arrow), whereas 4HR and insulin promote redistribution toward the cell periphery. Scale bars, 20 μm.

### 3.2 4HR activates AMPK–ACC signaling to transiently upregulate and stabilize GLUT4

qRT-PCR demonstrated that 4HR treatment (0.1, 1, and 10 μM) robustly upregulated GLUT4 mRNA expression at 2 h, with a 5–7-fold increase relative to control (Fig. 3a). Transcript levels declined toward baseline by 8 h, indicating rapid but transient transcriptional activation. Western blot analysis confirmed a corresponding transient increase in GLUT4 protein levels, peaking at 2 h and returning to baseline by 8-24 h (Fig. 3b).

**Figure 3.**
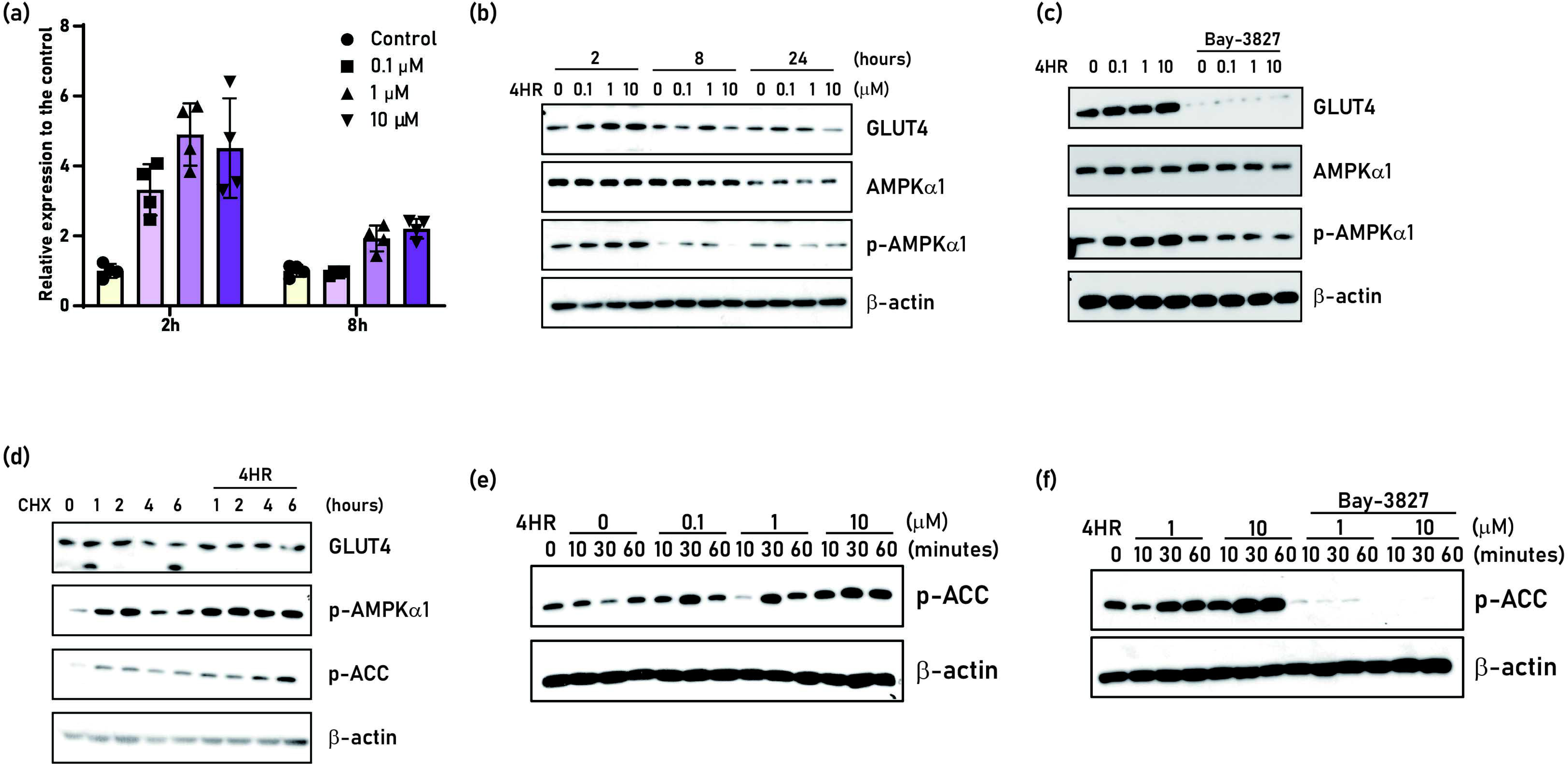
4HR activates the AMPK-ACC axis and increases GLUT4 expression via transcriptional upregulation and protein stabilization. (a) qRT-PCR of GLUT4 mRNA following 4HR treatment (0.1–10 µM) for 2 h and 8 h. Data are normalized to β-actin and expressed relative to vehicle (mean ± 95% CI). (b) Immunoblot analysis showing time- and dose-dependent effects of 4HR on GLUT4 protein, AMPKα1, and p-AMPKα1; β-actin as a loading control. (c) AMPK dependency: co-treatment with the AMPK inhibitor Bay-3827 for 2 h (?) attenuates 4HR-induced p-AMPKα1 activation and GLUT4 upregulation, while total AMPKα1 remains unchanged. (d) Cycloheximide (CHX) chase assay: in the presence of CHX, 4HR slows GLUT4 degradation over 0–6 h, coinciding with sustained p-AMPKα1 and p-ACC levels. (e) Short-term kinetics: 4HR increases p-ACC within 10–30 min in a dose-dependent manner. (f) Bay-3827 suppresses 4HR-mediated p-ACC induction, confirming AMPK dependency.

To test AMPK dependence, cells were co-treated with the AMPK inhibitor Bay-3827 for 2h. The inhibitor dose-dependently reduced 4HR–induced phosphorylation of AMPKα1 and attenuated GLUT4 protein expression, while total AMPKα1 levels remained largely unchanged (Fig. 3c), indicating a phosphorylation-specific mechanism. In a cycloheximide (CHX) chase assay, GLUT4 protein levels progressively declined in control cells, whereas 4HR treatment significantly slowed this decay over 0–6 h (Fig. 3d), suggesting enhanced protein stability. This stabilization coincided with sustained phosphorylation of AMPKα1 (p-AMPKα1) and ACC (p-ACC). Short-term exposure to 4HR (10-30 min) induced a dose-dependent increase in p-ACC levels (Fig. 3e), which was suppressed by Bay-3827, confirming AMPK dependence.

Treatment with the proteasome inhibitor MG132 markedly elevated GLUT4 levels, consistent with proteasomal degradation as a major turnover pathway (Fig. S1). Co-treatment with 4HR and MG132 did not further increase GLUT4 beyond MG132 alone, suggesting that 4HR stabilizes GLUT4 by attenuating proteasome-mediated degradation. Collectively, these findings indicate that 4HR modulates the AMPK–ACC axis to transiently enhance GLUT4 transcription and stabilize GLUT4 protein, thereby contributing to increased glucose uptake, potentially via CHC-sensitive trafficking.

### 3.3 4HR partially rescues body weight and muscle function in STZ-induced diabetic rats

Representative images at the end of the 8-week study revealed marked differences in body size among groups (Fig. 4a). STZ administration induced substantial weight loss compared with controls (p < 0.0001; Fig. 4b). Treatment with 4HR partially attenuated this weight loss, as the STZ/4HR group exhibited significantly higher terminal body weights than the STZ-only rats (p < 0.001). Neuromuscular function, assessed by Kondziela’s inverted-screen test (Fig. 4c), was significantly impaired by STZ-treated rats (p < 0.05). However, 4HR treatment improved performance, with STZ/4HR rats maintaining their grip duration significantly longer than

**Figure 4.**
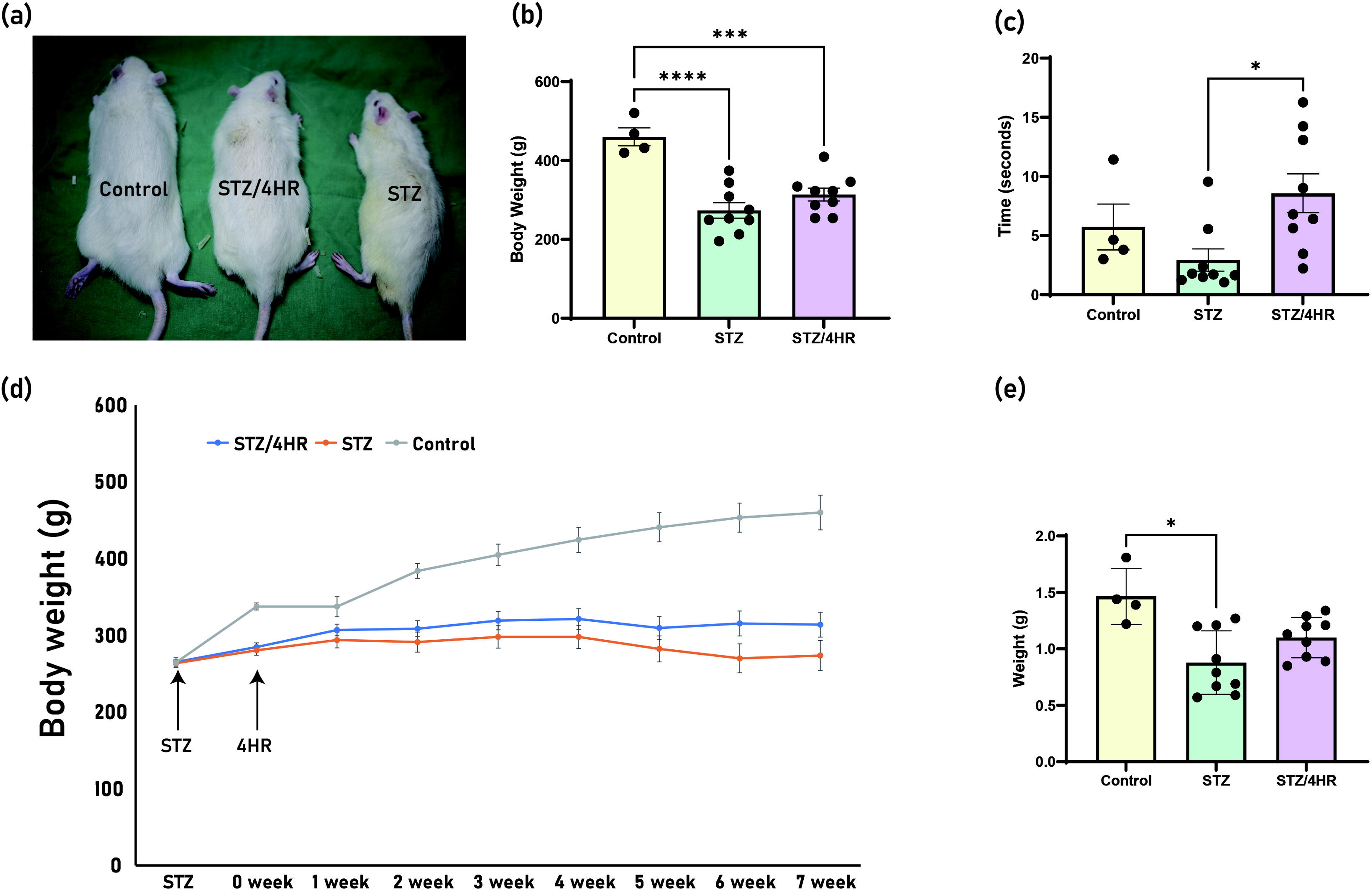
4HR partially restores body weight and neuromuscular function in STZ-induced diabetic rats. (a) Representative images of rats at study endpoint. (b) Terminal body weight (g) shows marked STZ-induced loss, partially rescued by 4HR (mean ± SD). (c) Kondziela’s inverted-screen test (time to fall, seconds): STZ impairs neuromuscular performance, improved by 4HR (mean ± SD). (d) Longitudinal body-weight trajectories (mean ± SEM). Arrows indicate STZ injection (week 0) and 4HR initiation (week 1). STZ/4HR rats consistently maintained higher weight than STZ rats from week 3 onward. (e) Gastrocnemius wet weight (g) reduced in STZ rats, partially preserved by 4HR (mean ± SD). Individual animals were shown as dots.

STZ rats (p < 0.05), suggesting a protective effect on muscle strength and endurance. Longitudinal monitoring of body weight (Fig. 4d) showed slow weight gain in both diabetic groups compared with controls. Following the initiation of 4HR treatment at week 1, the STZ/4HR trajectory diverged upward, consistently exceeding that of the untreated STZ rats from week 3 to week 7, demonstrating sustained partial rescue.

To determine whether these functional improvements were accompanied by preservation of skeletal muscle mass, gastrocnemius wet weight was quantified at sacrifice (Fig. 4e). Brown–Forsythe and Welch’s ANOVA indicated significant group differences (p = 0.0058 and p = 0.0208, respectively). Post hoc Dunnett’s T3 tests confirmed reduced gastrocnemius mass in STZ rats compared with controls (p = 0.0193), whereas the STZ/4HR group displayed intermediate values not significantly different from either group (p > 0.05). Collectively, these findings demonstrate that 4HR treatment partially counteracts STZ-induced reductions in body weight, skeletal muscle mass, and neuromuscular performance, suggesting a protective role in diabetic muscle health.

### 3.4 4HR lowers fasting blood glucose and preserves pancreatic islet integrity in STZ-induced diabetic rats

Fasting blood glucose was significantly elevated in STZ-treated rats compared with controls (*p* < 0.001; Fig. 5a). Administration of 4HR markedly reduced fasting glucose relative to the untreated STZ rats (*p* < 0.01), although levels remained above those of controls. In IPGTT, STZ rats exhibited persistently elevated glucose levels over 120 min with minimal decline from peak values (Fig. 5b). In contrast, 4HR-treated STZ rats demonstrated a modest but consistent reduction during the recovery phase, indicating partial improvement in glucose clearance.

**Figure 5.**
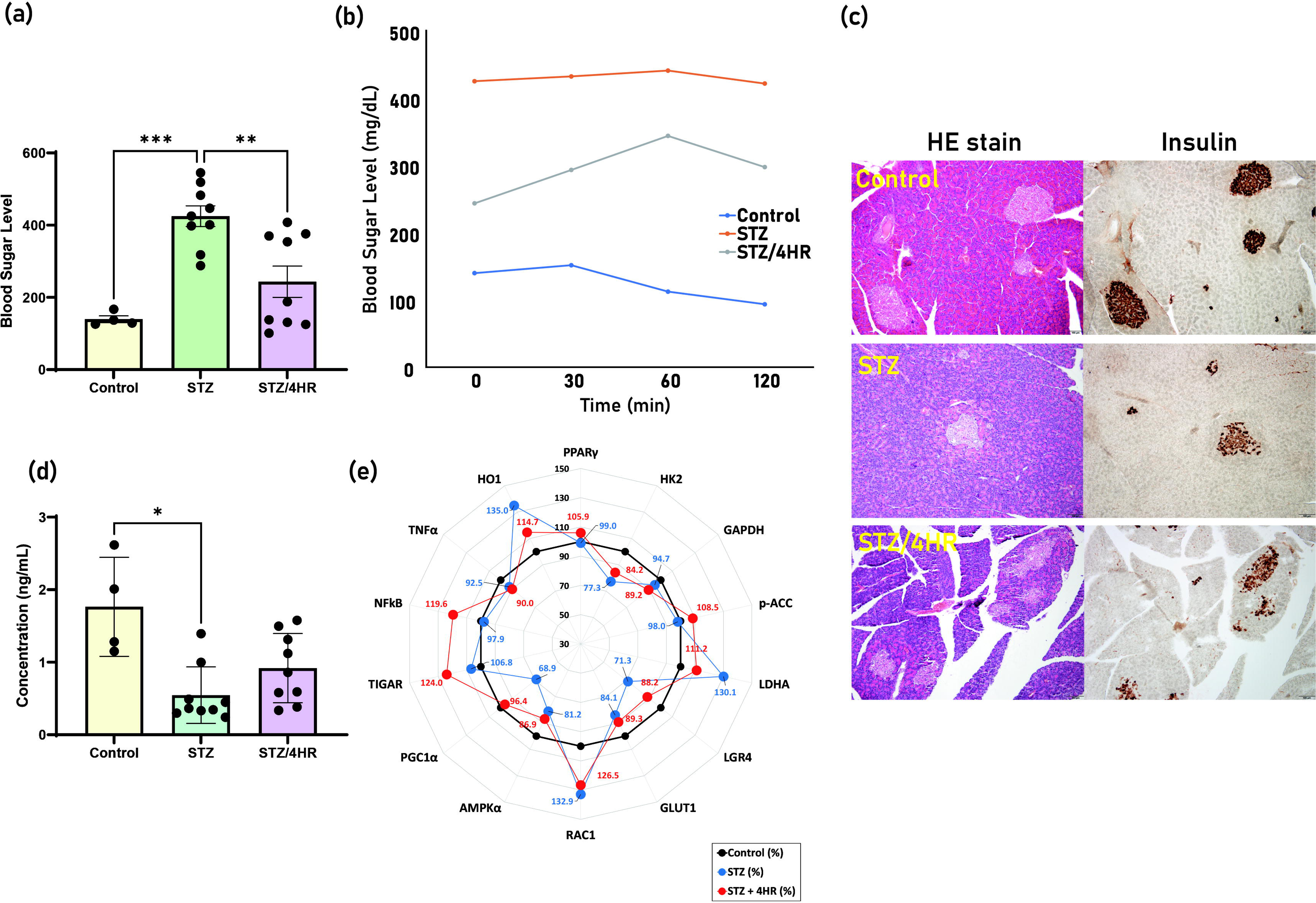
4HR improves glycemic control and modulates serum protein profiles in STZ-induced diabetic rats. (a) Fasting blood glucose: STZ markedly increased fasting glucose; 4HR reduced levels relative to STZ alone (mean ± SD). (b) IPGTT:.STZ rats showed persistently elevated glucose; STZ/4HR rats exhibited a modest improvement in clearance. (c) Pancreatic histology and insulin immunohistochemistry: STZ caused severe islet atrophy and insulin loss; 4HR preserved islet size and partially restored insulin staining. (d) Serum insulin concentration: reduced in STZ rats; 4HR showed a non-significant upward trend (mean ± SD). (e) Serum protein profiling via IP-HPLC: radar plot showing relative abundance (% of control) of HK2, GAPDH, LDHA, HO-1, LGR4, GLUT1, RAC1, PPARγ, p-ACC, TIGAR, PGC1α, AMPKα, NFκB, and TNFα.

Histological analysis of pancreatic tissue using H&E staining revealed severe islet atrophy and architectural disruption in STZ rats, whereas controls maintained normal islet morphology (Fig. 5c). 4HR treatment preserved islet structure with larger and more intact islets compared to the STZ rats. Immunohistochemistry for insulin demonstrated a marked reduction in β-cell insulin content in STZ rats, while 4HR partially restored insulin-positive staining. Serum insulin concentrations were significantly lower in STZ rats compared with controls (*p* < 0.05; Fig. 5d). Although 4HR did not significantly increase circulating insulin levels relative to STZ alone, a trend toward higher values was observed. These findings suggest that 4HR attenuates hyperglycemia and is associated with improved preservation of islet morphology in STZ-induced diabetes, although detailed quantitative analyses of β-cell mass and function were not performed.

### 3.5 4HR modulates serum protein expression profiles in STZ-induced diabetic rats

To avoid albumin-binding–related limitations inherent to serum Western blotting, including high background, weak signal intensity, and poor quantification, IP-HPLC was employed to profile serum proteins in control, STZ-diabetic rats (STZ-DR), and STZ-DR+4HR rats (Fig. 5e). The antibody panel targeted proteins involved in glycolysis, gluconeogenesis, and inflammation.

In STZ-DR, glycolytic enzymes HK2 and GAPDH were reduced to 77.3% and 94.7% of control levels, respectively. 4HR increased HK2 by 6.9% but slightly decreased GAPDH by 5.5%. LDHA and HO-1 were elevated in STZ-DR (130% and 135%) and reduced by 18.9% and 20.3% with 4HR. GLUT1 declined in STZ-DR (84.1%) and was modestly restored (+5.7%) by 4HR. LGR4, reduced to 71.3% in STZ-DR, increased by 16.9% with 4HR. RAC1 was elevated in STZ-DR (132.9%) and decreased by 6.4% after 4HR.

Among gluconeogenesis regulators, PPARγ remained near control levels (∼99%) in STZ-DR and increased by 6.9% with 4HR. p-ACC was slightly reduced (98%) in STZ-DR and increased by 10.5% after 4HR. TIGAR rose modestly in STZ-DR (106.8%) and further by 17.2% after 4HR. PGC1α was strongly suppressed in STZ-DR (68.9%) and increased by 27.5% following 4HR. AMPKα declined in STZ-DR (81.2%) and rose by 5.7% with 4HR. Regarding inflammatory mediators, NFκB expression was slightly reduced in STZ-DR (97.9%) and increased by 21.7% after 4HR, while TNFα was reduced in STZ-DR (92.5%) and remained largely unchanged (–2.5%) with 4HR.

Overall, 4HR attenuated glycolysis-promoting proteins elevated in STZ-DR, while upregulating gluconeogenesis regulators and partially restoring GLUT1 and LGR4. These molecular changes suggest improved metabolic regulation and cellular stress responses with 4HR; however, the exploratory nature of the serum profiling and the modest effect sizes warrant cautious interpretation, and causal links to inflammation or NFκB signaling remain speculative.

### 3.6 4HR restores glycogen content in skeletal muscle of STZ-induced diabetic rats

PAS staining of gastrocnemius muscle sections revealed abundant glycogen deposits in control rats, which were markedly diminished following STZ treatment (Fig. 6a). 4HR treatment substantially restored PAS staining intensity, indicating recovery of glycogen content. Quantitative analysis confirmed a significant increase in mean magenta intensity in the STZ/4HR group compared with the STZ-only rats (*p* < 0.05; Fig. 6b).

**Figure 6.**
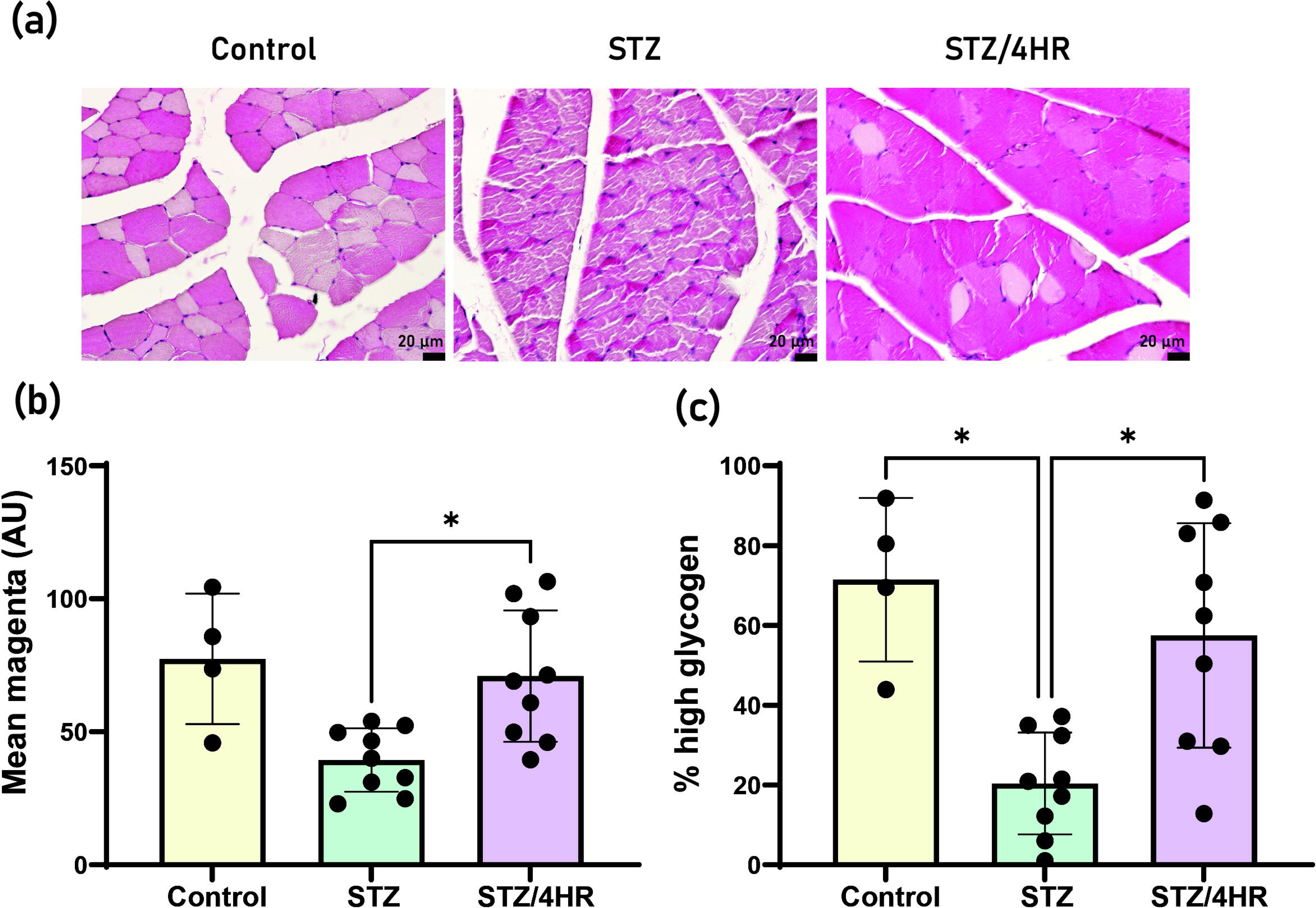
4HR restores skeletal-muscle glycogen content in STZ-induced diabetic rats. (a) Representative PAS-stained gastrocnemius sections: abundant glycogen in controls, reduced in STZ, restored with STZ/4HR. Scale bars:shown. (b) Quantification of PAS intensity (arbitrary units). 4HR significantly increased glycogen signal compared with STZ alone (mean ± SD). (c) Percentage of high-glycogen fibers per section, defined by threshold from controls. STZ reduced glycogen-rich fibers; 4HR significantly increased this fraction toward control levels. *p <0.05.

To assess glycogen distribution, muscle fibers were classified as “high-glycogen” based on a PAS-intensity threshold derived from controls. STZ markedly reduced the fraction of high-glycogen fibers, whereas 4HR treatment significantly increased this fraction relative to STZ (*p* < 0.05), approaching control levels (Fig. 6c). These findings indicate that 4HR treatment promotes a broader restoration of glycogen-rich fibers across skeletal muscle, supporting improved muscle energy storage in STZ-induced diabetes.

### 3.7 4HR reactivates AMPK–ACC signaling in diabetic skeletal muscle

Immunohistochemical analysis showed strong GLUT4 staining in the sarcolemma and cytoplasm of control muscle, which was nearly absent in STZ rats. 4HR treatment restored GLUT4 immunoreactivity to levels comparable to controls (Fig. 7a). In STZ rats, p-AMPK and p-ACC immunoreactivity in gastrocnemius fibers was markedly diminished, whereas 4HR administration restored both signals (Fig. 7b).

**Figure 7.**
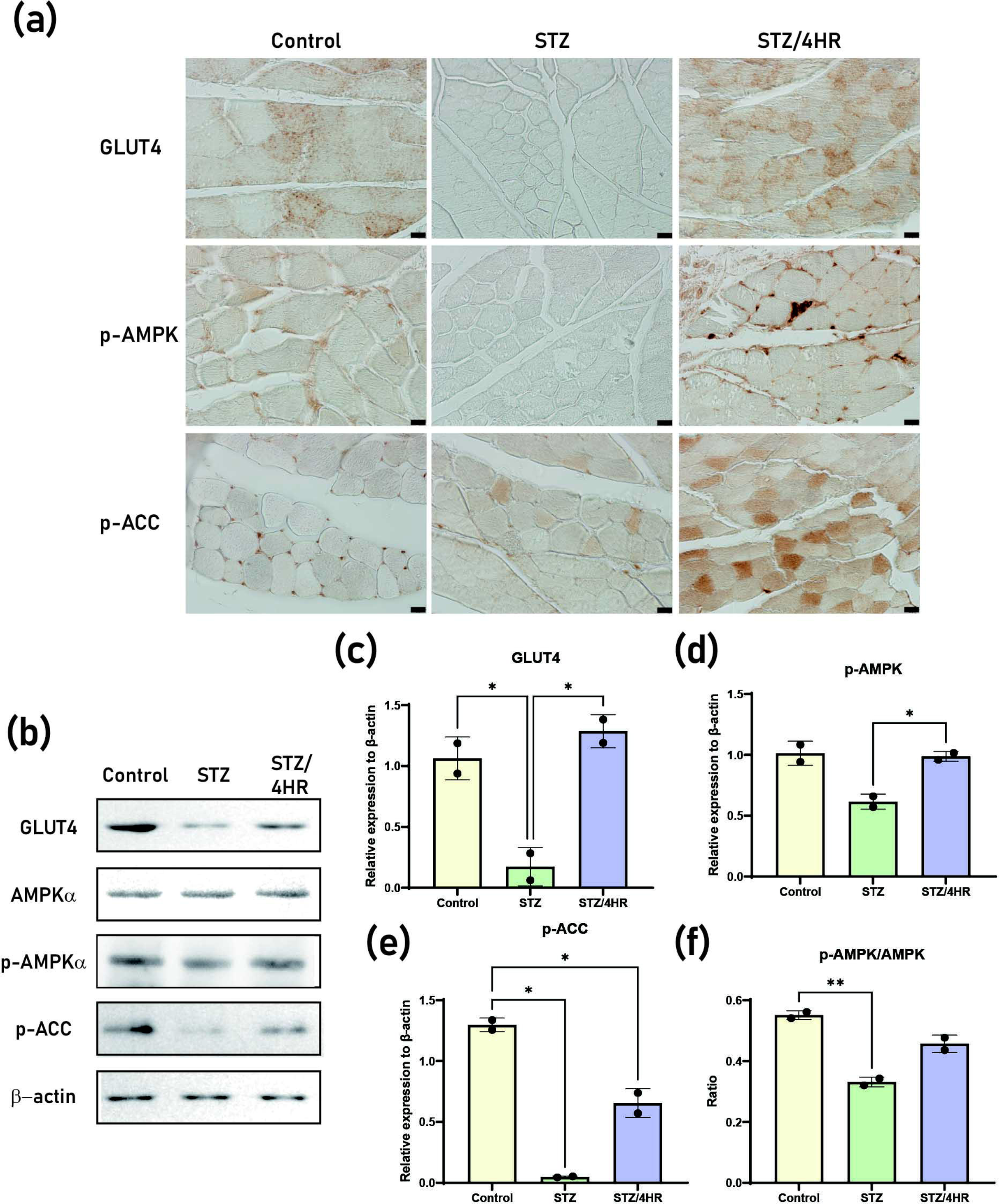
4HR restores GLUT4 expression and AMPK–ACC signaling in skeletal muscle. (a) Representative gastrocnemius muscle sections stained for GLUT4, p-AMPK, and p-ACC. Control muscle shows strong staining; STZ markedly reduces signals; 4HR restores toward control levels. Scale bars indicated. (b) Representative Western blots of gastrocnemius lysates probed for GLUT4, AMPKα, p-AMPKα (Thr172), p-ACC; β-actin as loading control. Quantitative densitometry:(c) GLUT4/β-actin, (d) p-AMPK/β-actin, (e) p-ACC/β-actin, and (f) p-AMPK/AMPKα ratio. Data are mean ± SD; individual animals shown as dots. *p<0.05, **p<0.01.

Western blot analysis corroborated the histological findings (Fig. 7b). GLUT4 protein was significantly reduced in STZ rats but restored in STZ/4HR rats (*p* < 0.05 vs STZ; Fig. 7c). Similarly, p-AMPKα and p-ACC levels were suppressed in STZ rats and significantly increased with 4HR treatment (Fig. 7d-e). The p-AMPKα/AMPKα ratio also improved with 4HR (*p* < 0.01 vs STZ; Fig. 7f). Collectively, these results demonstrate that 4HR reactivates AMPK-ACC signaling and restores GLUT4 expression in diabetic skeletal muscle.

## 4. Discussion

This study demonstrates that 4HR ameliorates multiple features of STZ-induced diabetic muscle dysfunction across molecular, histologic, functional, and metabolic levels. At the cellular level, 4HR rapidly induced phosphorylation of ACC, followed by sustained phosphorylation of AMPKα, accompanied by transient elevations in GLUT4 protein and slower degradation kinetics. These effects were attenuated by AMPK inhibition and partially resembled MG132 treatment, suggesting-but not conclusively proving-AMPK-dependent and proteostasis-related mechanisms (Fig. 2-3). *In vivo*, 4HR partially reduced fasting glycemia, mitigated STZ-induced weight loss, and improved neuromuscular performance (Fig. 4-5). In skeletal muscle, 4HR restored GLUT4 abundance and localization, reactivated the AMPK–ACC signaling, and preserved glycogen content (Fig. 6-7). Together, these findings support a model in which 4HR enhances muscle glucose handling by modulating GLUT4 expression, stability, and subcellular distribution, with downstream benefits for systemic metabolism and muscle function.

Consistent with canonical GLUT4 trafficking, 4HR promoted redistribution of GLUT4 from perinuclear pools toward peripheral regions. This pattern resembles early steps of GLUT4 vesicle mobilization, although confocal imaging cannot confirm true membrane insertion (Fig. 2). The observed redistribution parallels insulin- or contraction-induced mobilization of GLUT4 to the sarcolemma and T-tubule system [26]. Mechanistically, the temporal sequence aligns with AMPK-related pathways such as TBC1D1/TBC1D4 regulation [14], although these intermediates were not directly assessed. Loss of perinuclear puncta with concurrent peripheral enrichment supports this interpretation [27]. However, confocal redistribution alone may not be enough to confirm membrane fusion; Future studies using surface biotinylation or TIRF microscopy are warranted [28].

The timing of molecular events-p-ACC within 30 min, sustained p-AMPKα over 2–24 h, and a transient GLUT4 protein peak at 2 h—suggests an AMPK-first mechanism facilitating GLUT4 trafficking prior to net protein accumulation (Fig. 3). ACC phosphorylation serves as a sensitive readout of AMPK activation [29,30], and Bay-3827 suppression of both p-AMPK and GLUT4 supports this pathway. CHX-chase and MG132 experiments further suggest that 4HR influences post-translational turnover of GLUT4, potentially via proteasome-dependent degradation, although direct assays of ubiquitination or ERAD were not performed [31]. The non-additive effect of 4HR with MG132 implies partial overlap in proteasome-related pathways. In addition, 4HR induced a transient 5–7-fold increase in GLUT4 mRNA and slowed protein decay, consistent with coordinated transcription and post-translational regulation (Fig. 3). Previous reports describing 4HR as a pharmacologic chaperone [22,32], may provide one possible explanation, although this mechanism was not directly tested here.

In vivo, 4HR reduced fasting glucose and partially preserved body in STZ rats, despite only modest trends toward increased circulating insulin. These outcomes are compatible with improved peripheral glucose handling, likely involving AMPK-related pathways in skeletal muscle, together with partial preservation of pancreatic structure (Fig. 5). STZ-induced β-cell loss via GLUT2-mediated DNA alkylation, PARP activation, and oxidative damage explains the reduced insulin levels and islet atrophy [33] (Fig. 5). The intermediate phenotype of STZ/4HR rats aligns with prior reports of 4HR’s antioxidant and cytoprotective properties [21], suggesting partial preservation of β-cell integrity even without full recovery of insulin secretion.

Serum proteomic profiling revealed that 4HR downregulated glycolytic enzymes elevated in STZ rats while upregulating regulators of oxidative metabolism and insulin sensitivity (Fig. 5e). This metabolic shift suggests enhanced mitochondrial function and metabolic flexibility [34,35]. TIGAR elevation supports an antioxidant tilt via NADPH production [36]. Restoration of GLUT1 and LGR4 is favorable, given their role in basal glucose uptake and β-cell health [37]. Together, these findings suggest that 4HR attenuates oxidative stress-related β-cell injury.

PAS staining demonstrated glycogen recovery in skeletal muscle under 4HR treatment (Figure 6). In STZ diabetes, insulin deficiency impairs Akt→TBC1D4/AS160 signaling, limiting GLUT4 translocation, glucose entry, and glycogen synthesis [14]. Reduced glucose-6-phosphate (G6P) availability suppresses glycogen synthase activity [38]. Under 4HR, increased PAS intensity and restoration of high-glycogen fibers indicate widespread glycogen repletion. Elevated p-AMPK and p-ACC levels support an AMPK-first response (Fig. 7), enhancing glucose influx and G6P availability, thereby activating GS even under residual inhibitory phosphorylation [30,39]. Functionally, improved performance in the inverted-screen test aligns with restored glycogen stores, as glycogen depletion constrains muscle endurance and Ca²L handling [40].

We propose that 4HR mitigates diabetic sarcopenia through a multimodal mechanism centered on AMPK signaling: (i) rapid activation of AMPK and its downstream ACC phosphorylation shift cellular metabolism toward glucose uptake and oxidation while suppressing lipogenesis; (ii) transient GLUT 4 transcriptional upregulation and attenuated proteasomal degradation expand the functional GLUT4 reservoir; (iii) peripheral GLUT4 redistribution facilitates glycogen repletion and sustains ATP production during muscle contraction; and (iv) systemic improvements, including reduced fasting glucose, improved IPGTT recovery, partial preservation of islet integrity, and favorable serum protein remodeling, collectively mitigate catabolic drivers of muscle protein loss (Fig. 8). By simultaneously targeting substrate influx (GLUT4) and energy-sensing pathways (AMPK–ACC), 4HR addresses fundamental metabolic impairments in diabetic skeletal muscle that are insufficiently corrected by glycemic control alone.

**Figure 8.**
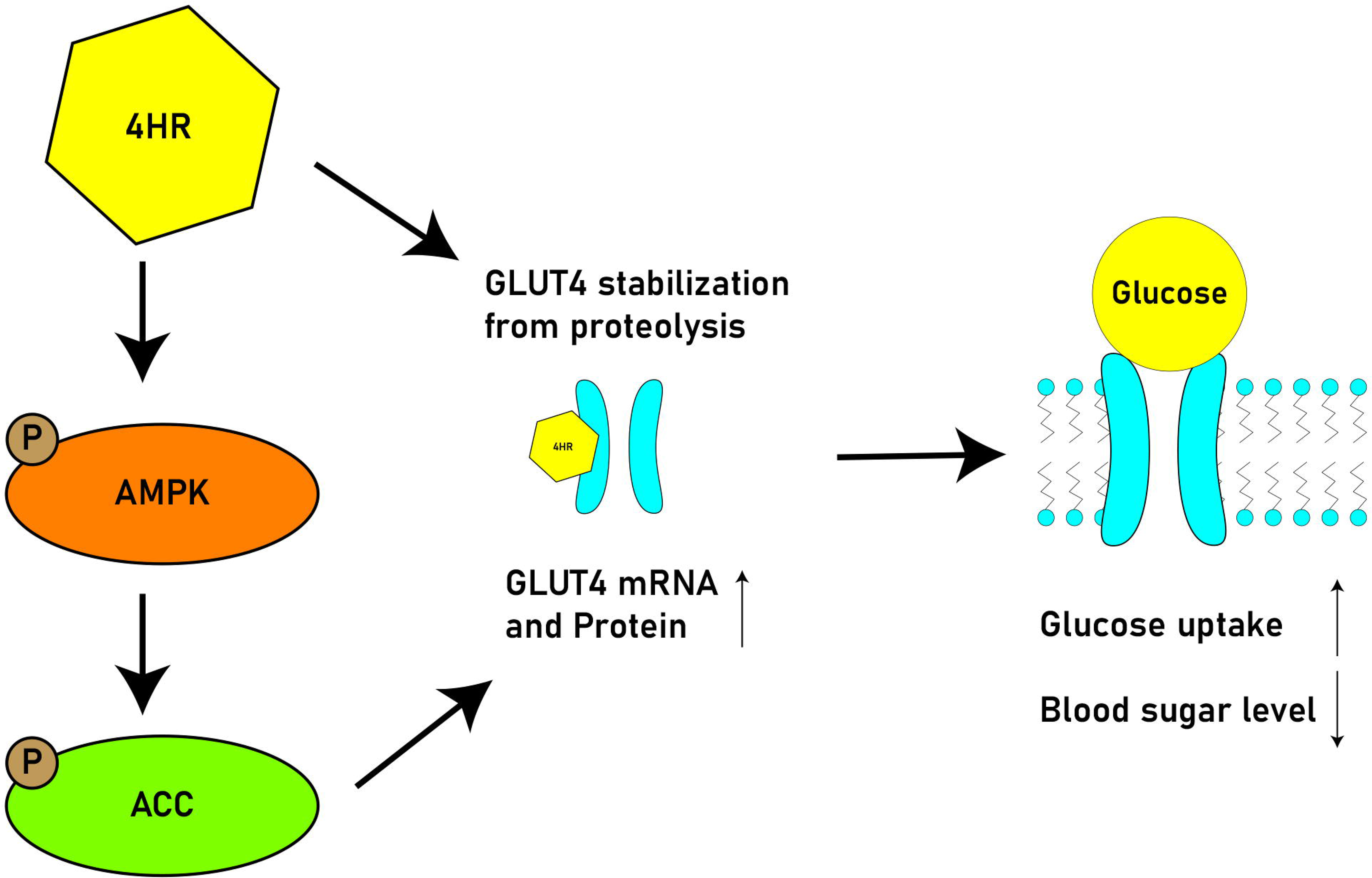
Proposed mechanistic model of 4HR in glucose regulation. 4HR activates AMPK, leading to phosphorylation of acetyl-CoA carboxylase (ACC) and enhanced downstream metabolic signaling. Concurrently, 4HR stabilizes GLUT4 by reducing proteolytic degradation and upregulates GLUT4 mRNA and protein expression. These combined effects facilitate GLUT4 translocation to the plasma membrane, thereby promoting glucose uptake and contributing to reduced circulating glucose levels.

Limitations include the absence of direct muscle glucose uptake assays, reliance on a single diabetic model, lack of pharmacokinetic profiling, and the exploratory nature of serum proteomics. Future studies should incorporate in vivo AMPK blockade, genetic models, and tissue-specific analyses to confirm the mechanistic necessity. Despite these constraints, consistent improvements across signaling pathways, transporter biology, glycogen storage, muscle function, and glycemic control support 4HR as a promising candidate for further preclinical evaluation in diabetic muscle dysfunction and metabolic disease.

## 5. Conclusion

4HR restored glucose homeostasis and improved muscle function in diabetic rats. In C2C12 myotubes, 4HR increased GLUT4 expression, altered subcellular localization, and enhanced glucose uptake in parallel with AMPK-related signaling changes. In STZ-induced diabetic rats, chronic 4HR administration reduced fasting glucose, improved glucose tolerance, preserved muscle glycogen, and ameliorated functional decline. These findings suggest that 4HR enhances skeletal muscle glucose handling in diabetes, although the precise mechanisms remain to be clarified. Given its safety profile and multifunctional activity, 4HR represents a promising candidate for further preclinical investigation in diabetic muscle dysfunction and metabolic disease.

## Acknowledgments

This work was carried out with the support of “Cooperative Research Program for Agriculture Science and Technology Development (Project No. RS-2024-00336327)” Rural Development Administration, Republic of Korea. This research was also funded by the National Research Foundation of Korea (NRF) grant funded by the South Korean Government (MSIT) (RS-2022-NR069290 and RS-2025-23963008)

## Ethical statement

All procedures were performed in accordance with the guidelines for laboratory animal care and were approved by the Gangneung-Wonju National University for animal research (GWNU-2024-24).

## Supplementary Materials

Detailed descriptions of the experimental procedures and information on the antibodies used in this study are provided in the Supplementary Materials.

## Authors’ contributions

S.-G.K. collected the data and drafted the manuscript. X.C. conducted cellular experiments. S.-K.H. and J.-H.O. carried out animal experiments. S.P., J.C., and S.-K.L. performed the blood analyses. D.W.K. conducted the Western blotting for tissue proteins. Y.Y.J. performed the histological analysis. J.-Y.C. and S.-G.K. revised the manuscript and prepared the figures. J.-Y.C. and S.-G.K. critically reviewed the final version of the article.

## References

1. American Diabetes Association. Diagnosis and classification of diabetes mellitus. Diabetes Care 2010; 33 Suppl. 1: S62–S69. 10.2337/dc10-S062

2. Chen H, Huang X, Dong M, Wen S, Zhou L, & Yuan X. The association between sarcopenia and diabetes: from pathophysiology mechanism to therapeutic strategy. Diabetes Metab Syndr Obes 2023; 16: 1541–1554. 10.2147/DMSO.S410834

3. Yuan S, & Larsson SC. Epidemiology of sarcopenia: prevalence, risk factors, and consequences. Metabolism 2023; 144: 155533. 10.1016/j.metabol.2023.155533

4. Jang HC. Sarcopenia, frailty, and diabetes in older adults. Diabetes Metab J 2016; 40: 182–189. 10.4093/dmj.2016.40.3.182

5. Ai Y, Xu R, & Liu L. The prevalence and risk factors of sarcopenia in patients with type 2 diabetes mellitus: a systematic review and meta-analysis. Diabetol Metab Syndr 2021; 13: 93. 10.1186/s13098-021-00707-7

6. Okamura T, et al. A multi-omics approach to overeating and inactivity-induced muscle atrophy in db/db mice. J Cachexia Sarcopenia Muscle 2024; 15: 2030–2045. 10.1002/jcsm.13550

7. Tremblay F, Lavigne C, Jacques H, & Marette A. Defective insulin-induced GLUT4 translocation in skeletal muscle of high fat-fed rats is associated with alterations in both Akt/protein kinase B and atypical protein kinase C (zeta/lambda) activities. Diabetes 2001; 50: 1901–1910. 10.2337/diabetes.50.8.1901

8. Sandri M, et al. Foxo transcription factors induce the atrophy-related ubiquitin ligase atrogin-1 and cause skeletal muscle atrophy. Cell 2004; 117: 399–412. 10.1016/S0092-8674(04)00400-3

9. Milan G, et al. Regulation of autophagy and the ubiquitin-proteasome system by the FoxO transcriptional network during muscle atrophy. Nat Commun 2015; 6: 6670. 10.1038/ncomms7670

10. Eshima H. Influence of obesity and type 2 diabetes on calcium handling by skeletal muscle: spotlight on the sarcoplasmic reticulum and mitochondria. Front Physiol 2021; 12: 758316. 10.3389/fphys.2021.758316

11. Hernández-Ochoa EO, Llanos P, & Lanner JT. The underlying mechanisms of diabetic myopathy. J Diabetes Res 2017; 2017: 7485738. 10.1155/2017/7485738

12. Jakobsen J, & Reske-Nielsen E. Diffuse muscle fiber atrophy in newly diagnosed diabetes. Clin Neuropathol 1986; 5: 73–77.

13. Kjøbsted R, et al. AMPK in skeletal muscle function and metabolism. FASEB J 2018; 32: 1741–1777. 10.1096/fj.201700442R

14. Sakamoto K, & Holman GD. Emerging role for AS160/TBC1D4 and TBC1D1 in the regulation of GLUT4 traffic. Am J Physiol Endocrinol Metab 2008; 295: E29–E37. 10.1152/ajpendo.90331.2008

15. Gaster B, & Hirsch IB. The effects of improved glycemic control on complications in type 2 diabetes. Arch Intern Med 1998; 158: 134–140. 10.1001/archinte.158.2.134

16. Cruz-Jentoft AJ, et al. Sarcopenia: revised European consensus on definition and diagnosis. Age Ageing 2019; 48: 16–31. 10.1093/ageing/afy169

17. Najm A, Niculescu AG, Grumezescu AM, & Beuran M. Emerging therapeutic strategies in sarcopenia: an updated review on pathogenesis and treatment advances. Int J Mol Sci 2024; 25: 4300. 10.3390/ijms25084300

18. Kim SG. Biomedical application of 4-Hexylresorcinol. Singapore: Springer; 2024. 10.1007/978-981-97-0637-2

19. Shariff R, et al. Superior even skin tone and anti-ageing benefit of a combination of 4-hexylresorcinol and niacinamide. Int J Cosmet Sci 2022; 44: 103–117. 10.1111/ics.12759

20. Che X, et al. 4-Hexylresorcinol enhances Glut4 expression and glucose homeostasis via AMPK activation and histone H3 acetylation. Int J Mol Sci 2024; 25: 12281. 10.3390/ijms252212281

21. Yen GC, Duh PD, & Lin CW. Effects of resveratrol and 4-hexylresorcinol on hydrogen peroxide-induced oxidative DNA damage in human lymphocytes. Free Radic Res 2003; 37: 509–514. 10.1080/1071576031000083099

22. Kim SG. 4-Hexylresorcinol: pharmacologic chaperone and its application for wound healing. Maxillofac Plast Reconstr Surg 2022; 44: 5. 10.1186/s40902-022-00334-w

23. Gaida D, Park YW, Kang YJ, & Kim SG. Therapeutic potential of 4-hexylresorcinol in reducing sarcopenia in diabetic masseter muscle. Maxillofac Plast Reconstr Surg 2025; 47: 2. 10.1186/s40902-025-00457-w

24. Yoon JH, Kim DW, Lee SK, & Kim SG. Effects of 4-hexylresorcinol administration on the submandibular glands in a growing rat model. Head Face Med 2022; 18: 16. 10.1186/s13005-022-00320-7

25. Kim MK, Yoon CS, Kim SG, Park YW, Lee SS, & Lee SK. Effects of 4-hexylresorcinol on protein expressions in RAW 264.7 cells as determined by immunoprecipitation high-performance liquid chromatography. Sci Rep 2019; 9: 3379. 10.1038/s41598-019-38946-4

26. Lauritzen HP, Galbo H, Toyoda T, & Goodyear LJ. Kinetics of contraction-induced GLUT4 translocation in skeletal muscle fibers from living mice. Diabetes 2010; 59: 2134–2144. 10.2337/db10-0233

27. Brumfield A, et al. Insulin-promoted mobilization of GLUT4 from a perinuclear storage site requires RAB10. Mol Biol Cell 2021; 32: 57–73. 10.1091/mbc.E20-06-0356

28. Koumanov F, Yang J, Jones AE, Hatanaka Y, & Holman GD. Cell-surface biotinylation of GLUT4 using bis-mannose photolabels. Biochem J 1998; 330: 1209–1215. 10.1042/bj3301209

29. Steinberg GR, & Carling D. AMP-activated protein kinase: the current landscape for drug development. Nat Rev Drug Discov 2019; 18: 527–551. 10.1038/s41573-019-0019-2

30. Herzig S, & Shaw RJ. AMPK: guardian of metabolism and mitochondrial homeostasis. Nat Rev Mol Cell Biol 2018; 19: 121–135. 10.1038/nrm.2017.95

31. Yu C, Cresswell J, Löffler MG, & Bogan JS. The glucose transporter 4-regulating protein TUG is essential for highly insulin-responsive glucose uptake in 3T3-L1 adipocytes. J Biol Chem 2007; 282: 7710–7722. 10.1074/jbc.M610824200

32. Kim YS, Kim DW, Kim SG, & Lee SK. 4-Hexylresorcinol-induced protein expression changes in human umbilical cord vein endothelial cells as determined by immunoprecipitation high-performance liquid chromatography. PLoS One 2020; 15: e0243975. 10.1371/journal.pone.0243975

33. Lenzen S. The mechanisms of alloxan- and streptozotocin-induced diabetes. Diabetologia 2008; 51: 216–226. 10.1007/s00125-007-0886-7

34. Chan MC, & Arany Z. The many roles of PGC-1α in muscle—recent developments. Metabolism 2014; 63: 441–451. 10.1016/j.metabol.2014.01.006

35. Bao W, et al. Plasma heme oxygenase-1 concentration is elevated in individuals with type 2 diabetes mellitus. PLoS One 2010; 5: e12371. 10.1371/journal.pone.0012371

36. Huang B, Lang X, & Li X. The role of TIGAR in nervous system diseases. Front Aging Neurosci 2022; 14: 1023161. 10.3389/fnagi.2022.1023161

37. McMillin SL, et al. Muscle-specific ablation of glucose transporter 1 (GLUT1) does not impair basal or overload-stimulated skeletal muscle glucose uptake. Biomolecules 2022; 12: 1734. 10.3390/biom12121734

38. Palm DC, Rohwer JM, & Hofmeyr JH. Regulation of glycogen synthase from mammalian skeletal muscle—a unifying view of allosteric and covalent regulation. FEBS J 2013; 280: 2–27. 10.1111/febs.12059

39. Marr L, et al. Mechanism of glycogen synthase inactivation and interaction with glycogenin. Nat Commun 2022; 13: 3372. 10.1038/s41467-022-31109-6

40. Ørtenblad N, Westerblad H, & Nielsen J. Muscle glycogen stores and fatigue. J Physiol 2013; 591: 4405–4413. 10.1113/jphysiol.2013.251629

