## Supplementary Materials for "4-Hexylresorcinol Enhances Skeletal Muscle Glucose Handling through the AMPK-GLUT4 axis in Diabetic Rats"

**Supplementary Methods**

**Cell Culture and Differentiation**

Mouse C2C12 myoblasts (ATCC CRL-1772) were cultured in high-glucose Dulbecco’s Modified Eagle’s Medium (DMEM; 4.5 g/L glucose) supplemented with 10% fetal bovine serum (FBS) and 1% penicillin–streptomycin (P/S). Cells were maintained at 37 °C in a humidified incubator with 5% CO₂. Cultures were passaged at 70–80% confluence, and passages 5–15 were used. For differentiation, myoblasts were grown to 80–90% confluence and switched to DMEM containing 2% horse serum (HS) and 1% P/S. Medium was changed every 48 h, and after 4–7 days elongated multinucleated myotubes were obtained. Experiments were performed on mature myotubes (day 5–6 of differentiation).

**Reagents**

4-Hexylresorcinol (4HR, ≥98% purity; Sigma-Aldrich, Cat# H5375) was prepared as a 100 mM DMSO stock and diluted into medium to final concentrations of 0.1–10 µM (final DMSO ≤0.1%). The AMPK inhibitor BAY-3827 (MedChemExpress, Cat# HY-112083) was dissolved in DMSO (10 mM stock) and used at 1–2 µM. The GLUT4 inhibitor, α-cyano-4-hydroxycinnamic acid (CHC; Sigma-Aldrich, Cat# 476870), was applied at 5 mM. The proteasome inhibitor MG132 (Sigma-Aldrich, Cat# M7449) was used at 10 µM. Cycloheximide (CHX; Sigma-Aldrich, Cat# C7698) was prepared at 100 mg/mL and applied at 50 µg/mL to inhibit protein synthesis. Vehicle controls contained equivalent DMSO.

**Glucose Uptake Assay**

Differentiated C2C12 myotubes were treated with 0–10 µM 4HR for 2 h. Glucose uptake was assessed using the fluorescent glucose analog 2-NBDG. Fluorescence was measured at 488/520 nm with a FlexStation 3 microplate reader.

**GLUT4 Translocation Assay**

For immunofluorescence, myotubes grown on coverslips were fixed with 4% paraformaldehyde and permeabilized (or left intact to detect surface GLUT4 only). Cells were incubated with anti-GLUT4 antibody (e.g., Abcam ab654, 1:100), followed by Alexa Fluor 488-conjugated secondary antibody. Nuclei were counterstained with DAPI, and images acquired by confocal microscopy. GLUT4 distribution was quantified by membrane-to-cytoplasm fluorescence ratios.

For surface biotinylation, intact cells were incubated with sulfo-NHS-biotin at 4 °C, quenched with glycine, and lysed in RIPA buffer. Biotinylated proteins were isolated with streptavidin beads and analyzed by Western blotting for GLUT4.

**Time-Course and Inhibition Studies**

Differentiated myotubes were treated with 0.1–10 µM 4HR for 8 or 24 h. Protein extracts were probed for GLUT4, AMPK, phospho-AMPK (Thr172), and phospho-ACC (Ser79). To test AMPK involvement, cells were pre-incubated with BAY-3827 for 30 min before 4HR addition. To assess protein stability, CHX chase assays were performed (0–6 h), with or without 4HR. For proteasome inhibition, cells were treated with MG132 (6 h) in the presence or absence of 4HR.

**Western Blot Analysis**

Cells were lysed in RIPA buffer with protease/phosphatase inhibitors. Equal protein (20–30 µg) was resolved by SDS-PAGE and transferred to PVDF membranes. Membranes were blocked and incubated with antibodies against GLUT4, AMPK, phospho-AMPK, phospho-ACC, β-actin, or GAPDH. HRP-conjugated secondaries were applied, and bands were detected by ECL. Densitometry was performed using SigmaScan Pro 5.0 (SPSS Inc. Chicago, IL, USA), normalizing to loading controls.

**qPCR Analysis**

RNA Extraction: Total RNA was extracted from C2C12 myotubes after treatments using TRIzol Reagent (Invitrogen, Carlsbad, CA, USA) or a column-based kit (e.g., Qiagen RNeasy Mini Kit) following the manufacturer’s protocol. For TRIzol, ca. 0.5–1 mL TRIzol was added per well of a 6-well plate, and lysates were collected and phase-separated with chloroform. The aqueous phase was then precipitated with isopropanol, washed with 75% ethanol, and RNA pellets were air-dried and resuspended in RNase-free water. Any genomic DNA contamination was removed by DNase I treatment (e.g., 0.1 U/µL for 15 min at 37 °C, followed by heat inactivation or phenol/chloroform cleanup). RNA yield and purity (A₂₆₀/A₂₈₀) were assessed with a NanoDrop spectrophotometer. Only samples with A₂₆₀/A₂₈₀ ≈ 1.8–2.0 were used for cDNA synthesis. Reverse Transcription (cDNA Synthesis): For each sample, 1 µg of total RNA was reverse-transcribed into complementary DNA (cDNA) using a high-capacity cDNA reverse transcription kit (Applied Biosystems, Foster City, CA, USA) or equivalent. The RT reaction (20 µL) contained 1 µg RNA, 4 µL of 5× RT buffer, 1 µL of 10 mM dNTP mix, 1 µL of MultiScribe reverse transcriptase, 1 µL of random hexamers (50 µM) or oligo(dT)₁₅ primers, and RNase-free water up to 20 µL. Reactions were incubated at 25 °C for 10 min (primer annealing), then 42 °C for 50 min (cDNA synthesis), and terminated by heating at 70 °C for 15 min. The resulting cDNA was diluted 1:5 with nuclease-free water for use in qPCR. Negative control reactions without reverse transcriptase were performed to ensure absence of genomic DNA amplification. Real-Time PCR (qPCR): Quantitative PCR was carried out on an ABI Prism 7500 Fast or Bio-Rad CFX96 real-time PCR system using SYBR Green chemistry. Each 10 µL qPCR reaction contained 5 µL of 2× SYBR Green master mix (e.g., Applied Biosystems PowerUp SYBR Green or Takara Premix Ex Taq), 0.4 µL of each forward and reverse primer (0.2 µM final concentration each), 1 µL of diluted cDNA (corresponding to 50 ng RNA input), and 3.2 µL of water. Thermocycling conditions were: initial activation at 95 °C for 2 min, followed by 40 cycles of 95 °C for 15 s (denaturation) and 60 °C for 30 s (annealing/extension). A melting curve analysis from 65 °C to 95 °C was performed at the end of cycling to confirm a single specific product for each primer pair (Supplementary Table 1). All primer sets were validated for efficiency (90–110%) by standard curve dilution and for specificity by single-peak melt curves. Data Analysis: Each sample’s Cₜ (threshold cycle) for target genes (GLUT4, AMPKα2) and the housekeeping gene (GAPDH) was determined in technical triplicate reactions. Replicates showing >0.5 Cₜ variance were re-run or excluded. The relative mRNA expression of target genes was calculated using the 2^–ΔΔCt^ method (Livak-Schmittgen). First, ΔCₜ = Cₜ_(target)_ – Cₜ_(housekeeping)_ was computed for each sample, then ΔΔCₜ = ΔCₜ_(treated)_ – ΔCₜ_(control)_. Fold-change in mRNA was expressed as 2^(-ΔΔCₜ)^, setting the vehicle control condition to 1.0. All qPCR experiments were performed on at least three independent biological samples, each measured in triplicate, and results were reported as relative expression (mean ± SEM) after normalization to the reference gene.

**Animal studies**

Male Sprague–Dawley rats (n = 22; 6 weeks old; 230–260 g on arrival; 270–290 g after 1-week acclimation) were housed under standard conditions with chow and water ad libitum. All procedures were approved by the GWNU IACUC (GWNU-2024-24). Four rats were retained as healthy controls (no STZ). The remaining 18 underwent a two-stage STZ protocol. After a 6–8 h fast, rats received STZ 50 mg/kg i.v. via tail vein (fresh in cold 0.1 M citrate buffer, pH 4.5). Three days later, tail-vein glucose was measured. Rats with >300 mg/dL were considered diabetic (first-stage responders, n=4) and received no second dose. Non-responders were given a second 50 mg/kg STZ; 14 additional rats then exceeded 300 mg/dL (second-stage responders). First- and second-stage responders were block-randomized within stage, yielding n=9 per diabetic group. Outcome assessors were blinded where feasible. After confirming diabetes, the STZ/4HR group received 4HR 10 mg/kg s.c., 5 days/week for 7 weeks; the STZ group received matched vehicle.

Body weight was recorded weekly. At 15 weeks of age, after a 8 h fast, fasting blood glucose was measured from the tail vein. For the intraperitoneal glucose tolerance test (IPGTT), a 50% (w/v) glucose solution in sterile saline was administered i.p. at 1 g/kg to all groups (Control, STZ, STZ/4HR). Blood glucose was measured at 0, 30, 60, and 120 min post-injection. Twenty-four hours before terminal sampling, forelimb strength/endurance was assessed using a wire-mesh grid (≈40–50 cm square, ~1 cm spacing) mounted on a frame 40–50 cm above a padded surface. Each rat was placed at the center, the grid was inverted to 180° within ~2 s, and latency to fall was recorded with a maximum cutoff of 20 s. Animals that did not fall within 20 s were assigned 20 s for that trial. Three trials were performed per rat with 1–2 min rest, and the mean latency was used for analysis. Testing was performed by an investigator blinded to group. Under deep enflurane anesthesia, cardiac blood was collected into heparinized tubes and centrifuged; plasma insulin was quantified by Rat Ins1 Insulin ELISA (Sigma-Aldrich RAB0904) per manufacturer instructions. Additional plasma was reserved for IP-HPLC. Animals were then euthanized by transcardial perfusion with 4% paraformaldehyde before tissue harvest.

**IP-HPLC**

Plasma was prepared from coagulated whole blood by centrifugation. Aliquots were subjected to immunoprecipitation with antisera against the following targets: hexokinase 2 (HK2), glyceraldehyde-3-phosphate dehydrogenase (GAPDH), lactate dehydrogenase A (LDHA), heme oxygenase-1 (HO-1), leucine-rich repeat-containing G-protein-coupled receptor 4 (LGR4), glucose transporter 1 (GLUT1), Ras-related C3 botulinum toxin substrate 1 (RAC1), tumor necrosis factor-alpha (TNF-α), nuclear factor kappa-B (NF-κB), AMP-activated protein kinase alpha catalytic subunit (AMPKα), peroxisome proliferator-activated receptor-gamma coactivator-1 alpha (PGC-1α), TP53-induced glycolysis and apoptosis regulator (TIGAR), phosphorylated acetyl-CoA carboxylase (p-ACC), and peroxisome proliferator-activated receptor-gamma (PPARγ). The immunocomplexes were analyzed by high-performance liquid chromatography (HPLC) to generate the IP-HPLC profiles. Protein expression was expressed as a percentage of the untreated control (=100%) and visualized as line graphs and radar (star) plots. The housekeeping control β-actin varied by ≤5% relative to untreated control and was used to confirm technical consistency across runs.

**Histological Analysis**

Gastrocnemius muscles and pancreas were fixed in neutral-buffered formalin, paraffin-embedded, and sectioned at 5 µm. Dewaxing and rehydration using a series of xylene and decreasing ethanol concentrations removed the paraffin wax from the sections. Hematoxylin staining was performed for 6 min, followed by a brief dip in 1% acid alcohol to remove excess hematoxylin. Subsequently, the slides were immersed in eosin solution for 1 min, staining the cytoplasm and extracellular matrix. After rinsing in tap water, the slides underwent dehydration again using an increasing ethanol series. Cleared with xylene or a xylene substitute, the slides were mounted with a glass coverslip and allowed to dry before examination under a light microscope.

For PAS staining, a commercially available kit (CAT#: ab150680, Abcam, Cambridge, UK) was used to detect glycogen, glycoproteins, and glycolipids in tissues. Following fixation, dehydration, clearing, embedding, sectioning, and deparaffinization, tissue sections were treated with a periodic acid solution to oxidize tissue components for 5 min. After incubation, slides were carefully rinsed with distilled water to remove residual acid and stained with Schiff’s reagent, which reacts with the oxidized tissue components for 15 min. The excess reagent was washed off with lukewarm tap water for 20 min, allowing full-color development. Counterstaining with hematoxylin for 1-min enhanced contrast between different cellular components. The slides were rinsed with tap water or immersed in a bluing solution to stain cell nuclei blue. The tissue sections were dehydrated using an ascending ethanol series to prepare the slides for microscopic examination. Following dehydration, the slides were cleared with xylene or a substitute, allowing the mounting medium and glass coverslip to adhere correctly. Additional copies of slides were stained without counterstaining them for the quantitative analysis. Bright-field images were captured on a research microscope with a 20× objective under identical illumination, white balance, and exposure settings across groups. For each animal, 3–5 non-overlapping fields from central gastrocnemius were imaged per section; the analyst was blinded to group. Images were exported as RGB TIFFs. Color deconvolution (H–PAS vector) was applied to isolate the PAS (magenta) channel. Background was excluded by thresholding the tissue area; the mean magenta intensity was measured per field. For measuring percentage of high-glycogen fibers, individual myofibers were delineated. The mean PAS intensity per fiber was extracted. A single, fixed threshold (T) was derived from the Control cohort and applied unchanged to all groups. Fibers with mean intensity ≥ T were classified as “high-glycogen.” The % high-glycogen fibers were calculated per section and averaged per animal.

**Immunohistochemistry**

Tissue sections (5 μm) were prepared for immunohistochemical analysis. After hydration, slides were treated with a peroxidase blocker for 7 min, washed, and followed by blocking with serum-free protein block (Agilent, Santa Clara, CA, USA) for 1 h. Primary antibodies (diluted 1:100) were applied to the slides and incubated overnight at 4 °C. After washing, slides were incubated with streptavidin-conjugated secondary antibody (Dako™ Real Envision, Agilent) for 30 min, followed by incubation with 3,3′-diaminobenzidine for colorization.

**Statistical Analysis**

All data are presented as mean ± standard error of the mean (SEM) from at least three independent experiments. Statistical analyses were performed using GraphPad Prism (v10.6, GraphPad Software, San Diego, CA) and SPSS (IBM, Armonk, NY) where appropriate. Prior to significance testing, data were checked for normal distribution (Shapiro-Wilk test) and homogeneity of variances (Levene’s test). For comparisons between two groups, an unpaired two-tailed Student’s t-test was used. For multiple-group comparisons, one-way ANOVA followed by Tukey’s post-hoc test was employed to determine statistically significant differences among groups (e.g., control vs 4HR, and 4HR vs 4HR+inhibitor conditions). In time-course experiments with multiple time points and treatments, two-way ANOVA was used to assess the effects of treatment, time, and their interaction, followed by Bonferroni post-hoc corrections. A p-value of <0.05 was considered statistically significant for all tests. Exact p-values are reported where relevant. Sample sizes (n) refer to biological replicates: for cell culture studies, n = number of independent wells or dishes from separate passages subjected to the same treatment. Each experiment was repeated at least three times on different days. Results in figures are annotated with significance indicators (p<0.05, *p<0.01, etc.) for treated vs control or as otherwise detailed in figure legends. All statistical analyses followed standard guidelines for biomedical research, ensuring that the conclusions are supported by appropriate tests.

**Figure S1. Proteasome inhibition.** MG132 elevates GLUT4 levels; co-treatment with 4HR does not further increase GLUT4, and CHX co-treatment suggests the effect is due to reduced proteasomal degradation rather than enhanced synthesis. For all immunoblots, treatment durations (min or h) and 4HR concentrations (µM) are indicated above each lane; β-actin is used as a loading control.

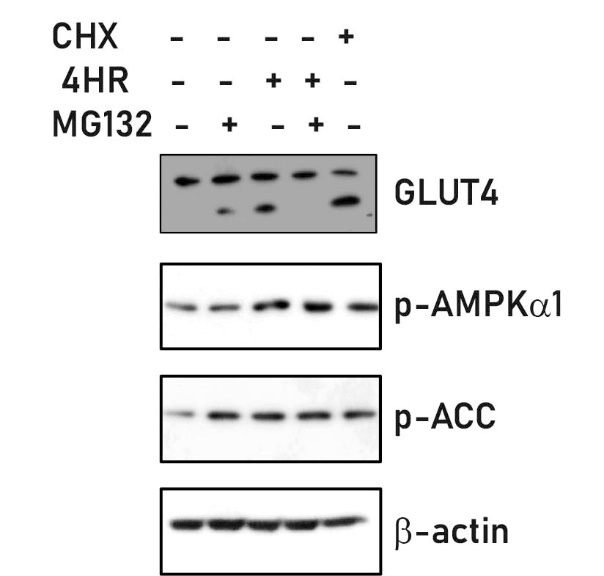

### **Table S1. Primer sequences used for quantitative RT-PCR**

| Gene | Forward primer (5′→3′) | Reverse primer (5′→3′) | Amplicon size (bp) |
| --- | --- | --- | --- |
| GLUT4 | AAAAGTGCCTGAAACCAGAG | TCACCTCCTGCTCTAAAAGG | ~150 |
| GAPDH | AGAACATCATCCCTGCATCC | CACATTGGGGGTAGGAACAC | ~120 |

Notes:
- Primer specificity was validated by melt curve analysis, showing single peaks.
- All primers amplified products between 100–200 bp.
- GAPDH was used as the endogenous control for normalization in the 2−ΔΔCt analysis.

### **Table S2. Antibodies used in this study**

| Target | Species | Type | Supplier | Cat# | Application | Dilution |
| --- | --- | --- | --- | --- | --- | --- |
| GLUT4 | Mouse | M | Santa Cruz Biotechnology | sc-53566 | WB, IP-HPLC, IF | 1:1000 |
| AMPKα1/2 (total) | Mouse | M | Santa Cruz Biotechnology | sc-74461 | WB, IP-HPLC | 1:1000 |
| Phospho-AMPKα (Thr172) | Rabbit | P | Cell Signaling Technology | #2535 | WB | 1:1000 |
| Phospho-ACC (Ser79) | Rabbit | P | Cell Signaling Technology | #3661 | WB | 1:1000 |
| HK2 | Rabbit | P | Cell Signaling Technology | #2867 | IP-HPLC | 1:500 |
| GAPDH | Rabbit | P | GeneTex | GTX627408 | WB, IP-HPLC | 1:2000 |
| LDHA | Rabbit | P | Cell Signaling Technology | #2012 | IP-HPLC | 1:500 |
| HO-1 | Rabbit | P | Abcam | ab13243 | IP-HPLC | 1:500 |
| LGR4 | Rabbit | P | Abcam | ab207623 | IP-HPLC | 1:500 |
| GLUT1 | Rabbit | P | Abcam | ab115730 | IP-HPLC | 1:500 |
| RAC1 | Mouse | M | BD Biosciences | #610650 | IP-HPLC | 1:500 |
| TNF-α | Goat | P | Santa Cruz Biotechnology | sc-1351 | IP-HPLC | 1:500 |
| NF-κB p65 | Rabbit | P | Cell Signaling Technology | #8242 | IP-HPLC | 1:500 |
| PGC-1α | Rabbit | P | Novus Biologicals | NB600-1415 | IP-HPLC | 1:500 |
| TIGAR | Rabbit | P | Abcam | ab37910 | IP-HPLC | 1:500 |
| PPARγ | Rabbit | P | Cell Signaling Technology | #2435 | IP-HPLC | 1:500 |
| β-actin | Mouse | M | Sigma-Aldrich | A5441 | WB | 1:5000 |

Notes:
- M: Monoclonal, P: Polyclonal.
